# Prolonged autophagy induction correlates with host cell protein reduction in CHO cell culture

**DOI:** 10.1101/2025.04.03.647032

**Authors:** Ansuman Sahoo, Taku Tsukiadate, Geetanjali Pendyala, Rong-Sheng Yang, Xuanwen Li, Sri Ranganayaki Madabhushi

**Author notes:** Equal contribution (Ansuman Sahoo, Taku Tsukiadate).

## Abstract

Autophagy, a cellular recycling process regulated by the CLEAR signaling pathway, plays a pivotal role in maintaining cellular homeostasis. We hypothesize that this process may regulate and reduce HCP levels by targeting intracellular proteins and organelles for degradation. This study investigates the relationship between autophagy induction and the reduction of high-risk host cell proteins (HCPs), including polysorbate-degrading enzymes (PSDEs), to enhance the stability of therapeutic biologics such as monoclonal antibodies (mAbs).

Using clonal analysis, we identified upregulation of the CLEAR pathway in one clone, correlating with a significant reduction in lipase activity and PSDE abundance. Furthermore, autophagy modulators, such as 3-methyladenine (3-MA), selectively decreased PSDE levels in both batch and fed-batch cultures. This resulted in 62% reduction in lipase activity that corresponded to a 22% improvement in polysorbate-80 stability. Additionally, 3-MA treatment increased mAb specific productivity and altered glycosylation profiles, increasing afucosylation and galactosylation levels. These findings highlight autophagy induction as a promising strategy to modulate product quality profiles and reduce high-risk HCPs in biologics production.

## Introduction

Recombinant proteins, particularly monoclonal antibodies (mAbs), have revolutionized therapeutic protein development in the biopharmaceutical industry, offering targeted treatments for various diseases. Over time, there has been an increasing demand for large-scale production of biologics, especially antibodies ^1^. Chinese hamster ovary (CHO) cells remain the preferred host system due to their robust growth in suspension cultures and ability to generate human-compatible glycosylation patterns ^2^.

To meet an increasing demand of biologics and the need to make biologics manufacturing more cost effective, strategies to improve productivity and product quality are cruicial. These have included strategies that increase specific productivity (Qp) of the recombinant cell lines through genetic engineering, subcloning, process optimization, or the use of enhancers such as sodium butyrate ^3, 4^. Recently, autophagy, a well conserved catabolic process involved in cellular homeostasis, has garnered significant attention for its role in modulating specific productivity. Autophagy helps degrade and recycle cellular components, including large protein complexes and organelles, through vacuolar engulfment in response to stressors ^5, 6^. While autophagy supports cell survival during conditions like starvation, growth factor deficiency, or hypoxia, excessive activation can result in autophagic cell death ^7^.

Rapamycin, an autophagy inducer, has been shown to enhance the survival of CHO cells in serum-free suspension medium ^8^. Apilimod, an FYVE finger-containing phosphoinositide kinase (PIKfyve) inhibitor significantly increased specific productivity (Qp) correlated with autophagy induction ^9^. Autophagy inducing peptide (AIP) has been shown to complete the autophagy process, which also boosts cell productivity ^10^. Similarly, application of 3-MA, SP600125, and dorsomorphin have a positive impact on Qp that have been proposed to induce autophagy rather than its inhibition ^11^. Although 3-MA has been traditionally described as a general inhibitor of autophagy, Baek et al. have demonstrated that it enhances autophagic flux rather than inhibiting the flux in CHO cells ^11^. This is consistent with the study by Wu et. al. who demonstrated autophagy induction in response to 3-MA under nutrient rich conditions ^12^. They have shown that 3-MA blocks class I PI3K activity persistently, the pathway known to inhibit autophagy. Beyond productivity, autophagy affects glycosylation patterns. For instance, autophagy inhibitors such as LY294002 decrease sialylation, whereas 3-MA has been reported to increase fucosylation, galactosylation, and sialylation ^13, 14^.

Apart from high productivity and desirable product quality profiles, ensuring host cell protein (HCP) clearance is critical for drug safety. HCPs are proteins released into the culture medium through secretion or cell lysis, and their presence in final drug formulations may compromise stability, induce immunogenicity, or degrade excipients ^15, 16^. Many of the “high-risk” HCPs exhibit reductase, glycosidase, or protease activity, which can lead to the breakdown of the product or formulation excipients, which in turn potentially promote aggregation in the drug formulation, thereby impacting its shelf life ^17^. For instance, lipases such as lipoprotein lipase (LPL), Sialate O-Acetylesterase (SIAE), and phospholipase A2 group VII (LP-PLA2), have the ability to degrade excipients in the final product formulations, specifically polysorbate 80 (PS-80) or polysorbate 20 (PS-20) ^18, 19^.

Upstream parameters can significantly influence the composition and level of HCPs, which can in turn impact the downstream separation process ^20, 21^. Addition of folic acid, glycine, and riboflavin supported an HCP reduction whereas MgCl_2_ increased the HCP concentrations in the Protein A eluted samples. Our previous research showed that culture parameters such as seeding density, and downshift temperature (TDS) have a substantial impact on the RNA level of many of the problematic HCPs ^22^.

However, the specific mechanisms by which signaling pathways and biological processes influence HCP production remain unclear. Autophagy and the CLEAR signaling pathway are closely interconnected. The transcription factor EB (TFEB), a master regulator of the CLEAR pathway, drives the expression of autophagy and lysosomal genes, enhancing autophagosome formation and lysosomal degradation under stress or starvation conditions ^23^. We hypothesized that these processes may play a role in the regulation and reduction of HCP levels, particularly by targeting and degrading intracellular proteins and organelles.

In this work, we leveraged insights from a clonal comparison of CHO cells producing a mAb to explore the correlation between the autophagy, lipase activity, and HCP abundance in the culture medium as well as Protein A purified samples. Our findings reveal that upregulation of autophagy and the CLEAR pathway through specific chemicals correlates with significant reductions in lipase activity and associated high-risk HCP abundance. This study highlights the potential of autophagy modulation as a strategy to enhance productivity and product quality by selectively reducing polysorbate-degrading enzymes (PSDEs), thereby improving the stability and efficacy of monoclonal antibodies.

## Materials and Methods

### 2.1 Cell Line

Two clones, each expressing the same monoclonal antibody (IgG mAb), were used in this study. These clones were derived from a glutamine synthetase (GS) knockout CHO cell line. Cells were maintained in a chemically defined CHO medium supplemented with 12.5 µM methionine sulfoximine (MSX, Sigma-Aldrich) and cultured in shake flasks within a humidified Multitron incubator at 36.5°C, 5% CO, and 140 rpm.

### 2.2 Cell Culture Operations

For fed-batch experiments with 3-methyladenine (3-MA), clones were propagated in shake flasks before being transferred to AMBR250 bioreactors (Sartorius Stedim, Göttingen, Germany). Bioreactors were equipped with two pitched-blade impellers and an open pipe sparger. Cells in the exponential growth phase were inoculated at a density of 0.5 × 10 cells/mL in 190 mL of Dynamis® culture medium (Gibco, NY, USA). From day 3, two proprietary feeds (Feed A and Feed B, 9:1 ratio) were added daily at 3% v/v. Glucose levels were maintained at 6 g/L by supplementation when they fell below 3 g/L. On day 5 or at a cell density of 15 × 10 cells/mL, the bioreactor temperature was reduced from 36.5°C to 33°C. The pH was maintained at 7.2 using sodium carbonate when it dropped below 6.9.

For batch cultures, Clone A was cultured in 24-well plates for seven days at 300 rpm, with glucose supplementation when levels fell below 3 g/L. 3-MA was added on day 3 at a final concentration of 5 mM, while autophagy-inducing peptide (AIP) was added at 2 µM on day 0. At least three biological replicates were performed for each treatment condition.

### 2.2 Cell Culture Sample Analysis and Analytical Methods

Daily cell culture samples were collected for immediate analysis. Viable cell density (VCD) and cell viability were measured using the trypan blue exclusion method with a Cedex HiRes cell counter (Roche Diagnostics GmbH, Mannheim, Germany). Concentrations of glucose, lactate, glutamine, and glutamate were determined with a Flex2 analyzer (Nova Biomedical). Antibody titer (g/L) was analyzed using affinity (HPLC) based Protein A column. Total lipase activity was determined with our proprietary assay involving FRET (Fluorescence Resonance Energy Transfer) based molecular probe.

### 2.3 Protein A affinity chromatography

Protein A affinity chromatography was carried out using MabSelect SuRe (Cytiva) resin, which was packed into a column with a total volume of 6.74 mL and a bed height of 19.7 cm. The residence time during the experiments was set to 4 minutes. An ÄKTA™ avant 25 system was employed for these chromatographic runs, with approximately 30 g/L of HCCF loaded onto the column for each experiment. Prior to loading, the column was sanitized with 5 column volumes (CVs) of sanitization buffer (0.1 M NaOH), followed by equilibration with 5 CVs of equilibration buffer (10 mM NaPO4, pH 6.5). After loading the HCCF, three wash steps were performed, each with 3 CVs of the respective wash buffers. Wash 1 and Wash 3 were conducted using the equilibration buffer, while Wash 2 involved 10 mM NaPO4 and 0.5 M NaCl (pH 6.5). For the elution step, 5 CVs of elution buffer (20 mM CH3COONa, pH 3.7) were applied, and the protein A product (PAP) was collected. Following elution, the column was stripped using 3 CVs of strip buffer (100 mM CH3COOH).

### 2.3 Proteomics Sample preparation and Analysis

The SP3 (Single-Pot Solid-Phase-enhanced Sample Preparation) method was used to process the cell culture fluid (HCCF) as described by previous studies ^24^. In brief, 20 µL of HCCF was incubated in a solution containing 4 M urea, 25 mM TCEP-HCl, and 50 mM acrylamide in 50 mM tris-HCl (pH 8.0) to a final volume of 50 µL at 60 °C for 30 minutes. Subsequently, an aliquot of 20 µL was combined with 10 µL of Sera-Mag SpeedBeads (200 µg; GE Healthcare) and 70 µL of acetonitrile, followed by incubation at 20 °C for 10 minutes while shaking at 500 rpm. The beads were washed three times with 200 µL of 80% ethanol. For protein digestion, the beads were treated with 20 µL of 50 mM tris-HCl (pH 8.0) containing 0.4 µg of trypsin and 0.1 µg of LysC at 37 °C for 18 hours. The peptide concentration was quantified using either the Nanodrop A280 or the BCA assay (Thermo Scientific) and adjusted to 0.02 g/L using 0.1% formic acid. Finally, 20 µL of the digested peptides (400 ng) was loaded onto an Evotip Pure (Evosep) for further analysis, following the manufacturer’s guidelines.

DIA (data independent acquisition) -based quantitative proteomics was performed on the Evosep One LC-Bruker timsTOF Pro 2 system in diaPASEF (parallel accumulation–serial fragmentation) mode ^25^. Peptides were separated using a Pepsep C18 column (8 cm × 100 μm, 3 μm) with gradient time of 11.5 min. The Bruker timsTOF Pro 2 system operated with 100 ms TIMS accumulation and ramp times, covering ion mobility ranges of 0.7–1.4 1/K0 and mass ranges of 300–1200 Th. Spectral libraries were generated from pooled sample measurements using ion mobility–coupled gas phase fractionation (IM-GPF), with the appropriate sample-specific methods and injection parameters. For IM-GPF, the TIMS ramp time was set to 100 ms, and the mass windows were adjusted to capture optimal resolution for peptide identification. Proteomics data were processed with DIA-NN v1.8.1, setting mass accuracies to 10 ppm for spectral library generation and 15 ppm for subsequent processing, with MBR disabled. For in-silico spectral library generation, precursor charge was restricted to 2−4 and the mass range to 300–1200 Th. The false discovery rate (FDR) thresholds for precursor and protein identification were set to 1%. An in-house sequence database, including the CHO proteome, proprietary product sequences, and common contaminants, was used for all analyses.

### 2.4 Statistical analysis

Differential expression analysis was conducted using the limma R package ^26^, fitting gene-wise linear models with lmFit, and comparing clones at each timepoint with makeContrasts and contrasts.fit. Moderated t-statistics were computed using eBayes. P-values were extracted with topTable, combined across timepoints using cumulative fold-changes, and adjusted with the Benjamini-Hochberg procedure. Combined results informed GSEA analysis ^27^.

### 2.5 Characterization of PS-80 degradation by MAX-UPLC-ESI-HRMS

Experiments were conducted using a Waters Acquity UPLC Classic connected to a Xevo G2-XS time-of-flight (TOF) mass spectrometer (Waters) equipped with an ESI source. Mixed-mode chromatography was performed on an Oasis MAX Column (2.1 x 20 mm, 80Å, 30 µm, Waters). The mobile phase initially consisted of 99% solvent A (0.1% trifluoroacetic acid in water) and 1% solvent B (0.1% trifluoroacetic acid in acetonitrile), held for 3 minutes. The gradient was then increased as follows: to 20% B at 3.1 min, to 25% at 7 min, to 80% at 10.1 min (held for 5 min), to 99% at 15.1 min (held for 4 min), and back to 1% at 19.1 min (held for 3 min). The flow rate was set at 0.4 mL/min, with a column temperature of 50°C. A 5 μL sample (1 μg of PS80) was injected for analysis.

Mass spectrometry parameters were as follows: capillary voltage at 3.0 kV, sampling cone voltage at 40 V, collision voltage at 10 V, source temperature at 150°C, desolvation temperature at 500°C, cone gas flow at 30 L/h, and desolvation gas flow at 850 L/h. Full-scan ESI mass spectra were collected over the m/z range of 120–4000 in resolution mode. Data acquisition was performed using Waters MassLynx 4.2 software. Waters raw data files were imported into Expressionist 18.0 for processing without prior conversion. The analysis included recalibration of m/z values using background ions, retention time (RT) restriction (maximum RT of 20 minutes), chemical noise removal with chromatogram smoothing, and specific settings for peak detection and isotope clustering (RT tolerance of 0.05 min, m/z tolerance of 25 ppm).

### 2.6 Western Blotting

Protein samples were generated using mammalian lysis buffer (ab179835, Abcam, USA) containing Halt Protease and phosphatase inhibitor cocktail (Catalog number 78440, Thermo Fisher Scientific, USA) following manufacturer instructions. Protein concentrations were determined by nanodrop based A280 measurement and 3 µg of protein lysates were separated by SDS-PAGE precast gel (Bolt Bis-Tris 4-12 %, Invitrogen, USA). Proteins were transferred to PVDF membrane using iBlot system (Thermo Fisher, Waltham, MA, USA). Antibody incubation, blocking, and washes were performed using the automated iBind Western System (Invitrogen, USA) according to the manufacturer’s instructions. The primary antibodies used were as follows: p62/SQSTM1 Antibody (NBP149954, Novus Biologicals); LC3B Polyclonal Antibody (Thermo Scientific PA146286, Invitrogen); beta-Actin Antibody (Rabbit polyclonal, Cell Signaling Technology 4970T); Phospho-Rb (Ser807/811) rabbit Antibody (Cell Signaling Technology, 8516T); TFEB antibody (Cell Signaling Technology, 4240S); LAMP1 (D2D11) Antibody (Cell Signaling Technology, 9091T); Caspase-7 Antibody (Cell Signaling Technology, 9492T). The detection was performed with HRP conjugated goat anti-rabbit IgG (Novus Biologicals, NB7160). The dilutions for primary antibodies were 1:1000 and that for secondary antibody was 1:500. The blots were imaged with ChemiDoc (BioRad) and intensity levels were quantified with BioRad Image Lab software. The intensity values were normalized against the intensity of beta-Actin.

## Results and Discussion

### 3.1 Upregulation of the CLEAR Signaling Pathway Correlates with Reduced Lipase Activity and HCP Abundance

We cultured these two clones in a 14-day fed batch culture in shake flask. The process parameters are included in a related paper (Figure 1, supplementary figure S1) ^28^. Both clones exhibited comparable overall process performance. Clone A displayed higher glutamine catabolism, indicated by lower glutamine levels and elevated ammonia accumulation. Additionally, Clone A maintained slightly higher viability and viable cell density, while titer and product quality profiles remained similar between the two clones. To investigate potential differences in host cell protein (HCP) composition, we conducted data-independent acquisition (DIA)-based quantitative proteomics on the cell culture fluid and identified 1961 HCPs. Notably, Clone A exhibited higher lipase activity (Figure 1A) and an increased abundance of high-risk HCPs, as previously outlined by Jones et al.^29^ and Park et al.^21^ (Figure 1B). Of importance, many lipases and polysorbate degrading enzymes (PSDEs) such as LIPA, PLA2G7, PLA2G15, PPT1 and PPT2 were more abundant in Clone A.

**Figure1:**
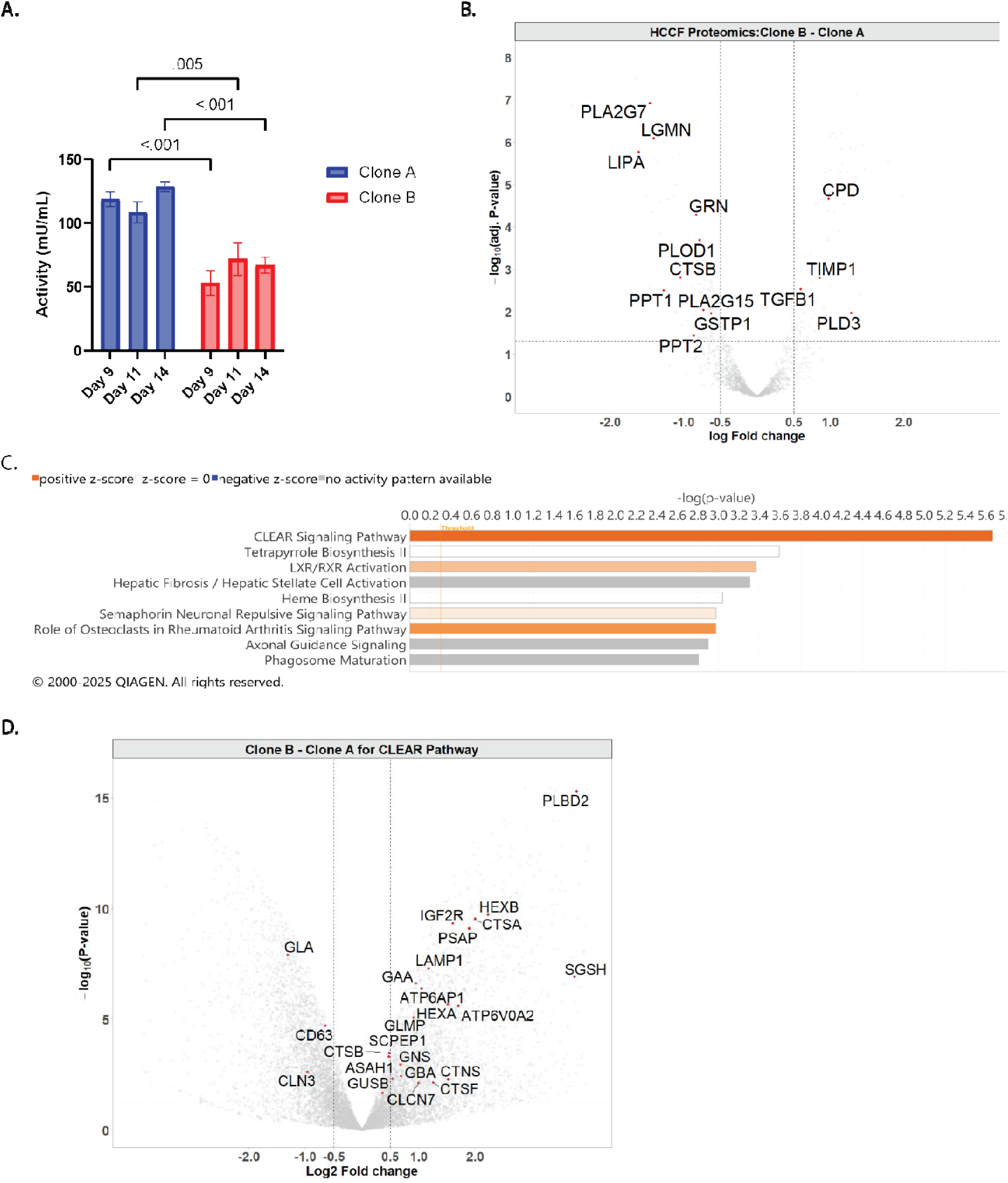
Molecular and functional comparisons between Clone A and Clone B. (A) Lipase activity measurements for Clone A and Clone B at multiple time points (Day 9, Day 11, and Day 14). (B) Proteomic analysis of cell culture fluid (HCCF) showing differences in host cell protein (HCP) profiles between Clone A and Clone B. (C) Pathway activity analysis of differentially expressed genes between clones, showing top enriched pathways. The orange -colored bars indicate predicted pathway activation and the gray bars indicate no activity prediction (D) Volcano plot of proteomic differences between Clone B and Clone A, with CLEAR pathway-related genes highlighted. Error bars represent the standard error of mean. Student’s t-test. ns: not significant; *: p < 0.05.

To further understand the molecular basis of these differences, we analyzed previously performed bulk RNA sequencing (RNA-seq) on Clone A and Clone B at the 250 mL bioreactor scale ^28^. Differential gene expression analysis from day-14 transcriptomics was processed using Qiagen IPA®, which identified the CLEAR (Coordinated Lysosomal Expression and Regulation) pathway as the most significantly activated pathway in Clone B (Figure 1C). This pathway, regulated by TFEB, promotes lysosomal biogenesis and autophagy by driving the transcription of genes containing CLEAR motifs in response to cellular stress or nutrient deprivation ^23^. Consistently, we observed upregulation of TFEB-regulated genes in Clone B, including lysosomal hydrolases (Ctsa, Ctsb, Hexa, Gaa, Scpep1, and Sgsh), as well as lysosomal membrane markers (Lamp1) and genes involved in lysosomal acidification (Atp6v0a2) (Figure 1D). These findings strongly suggest enhanced activation of the CLEAR pathway in Clone B, leading to increased lysosomal catabolic activity.

In summary, the upregulation of the CLEAR pathway in Clone B, which enhances autophagy and lysosomal catabolic processes, may contribute to the intracellular sequestration of lipases, thereby reducing their extracellular release and maintaining lower lipase activity. Based on these observations, we hypothesized that modulating autophagy could influence lipase and PSDE levels in the cell culture fluid.

### 3.2 Selective reduction of PSDE levels in batch culture HCCF with 3-MA/AIP

To test the hypothesis that autophagy induction reduces lipase levels, we evaluated the effects of autophagy-inducing agents including autophagy inducing peptide (AIP)^10^ and 3-methyladenine (3-MA). To test the effect of these compounds, we first performed a 7-day batch study in the presence of these compounds using clone A with the goal of reducing the lipase abundance. Figure 2A shows that the process performance of culture treated with 3-MA (5 mM) as well as AIP (2 µM). Overall, the trends were comparable except that the 3-MA treated culture had lower IVCD (integrated viable cell density) and higher Qp (specific productivity) while maintaining similar titer when compared to control conditions. This effect of 3-MA was consistent with previous reports ^30^. In contrast, 2 µM AIP ^10^ supplementation did not significantly impact Qp and titer.

**Figure 2:**
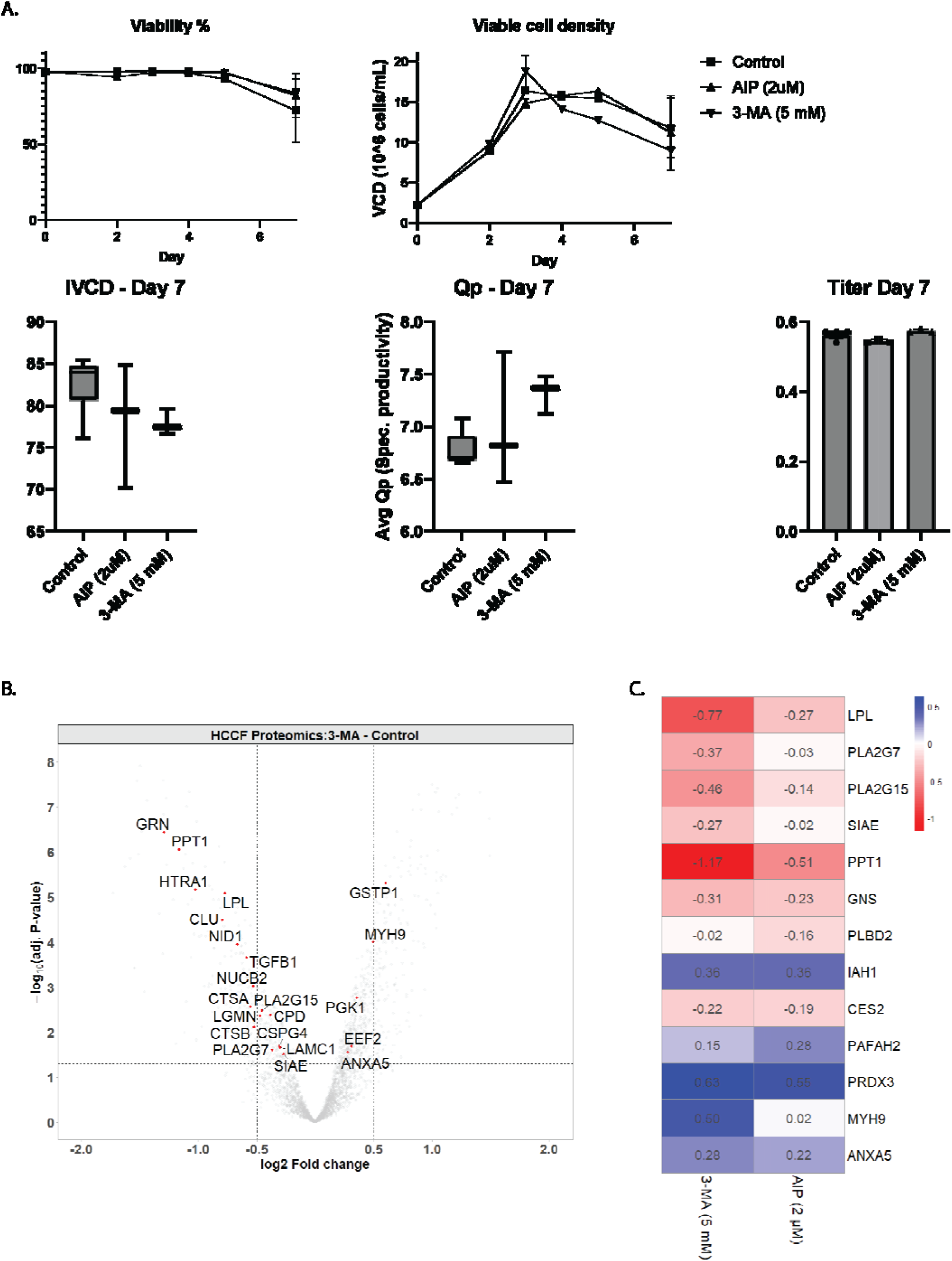
Evaluation of 3-MA effects on cell culture performance and HCP profile in batch culture. (A) Cell culture performance metrics, including viability (%), viable cell density (VCD, ×10 cells/mL), integral viable cell density (IVCD), specific productivity (Qp), and titer on Day 7 under control, AIP (2 µM), and 3-MA (5 mM) conditions. (B) Volcano plot illustrating the proteomic differences in host cell culture fluid (HCCF) between the control and 3-MA-treated cultures, highlighting the reduction in problematic HCPs with 3-MA treatment. (C) Heatmap showing the abundance differences relative to the control for select HCPs under 3-MA and AIP treatments, highlighting the similarities between the two treatments.

Proteomics analysis of the cell culture supernatant revealed differences in HCP abundance across treatment conditions. As shown in Figure 2B, 3-MA treatment led to a significant reduction in several lipases and PSDEs compared to the control. Figure 2C presents a heatmap highlighting key high-risk lipases (LPL, PLA2G7, PLA2G15), other polysorbate-degrading enzymes (SIAE, LIPA, PPT1), and additional high-risk HCPs. While AIP-treated cultures exhibited similar trends, the observed changes were less pronounced and did not reach statistical significance. Despite the smaller effect size, the overall pattern of HCP reduction in AIP-treated cultures mirrored that of 3-MA, suggesting that an optimized AIP concentration may be required for a more robust response. Taken together, these findings demonstrate that autophagy-inducing agents can effectively lower lipase and PSDE levels in cell culture fluid.

### 3.3 Selective reduction of PSDE activity in fed-batch culture supplemented with 3-MA

Fed-batch is the most commonly used process mode for biologics manufacturng. Given the promising reduction in HCPs using 3-MA in batch experiment, we tested the impact of 3-MA supplementation in a fed-batch process. Clone A was inoculated in ambr®250 bioreactors at a seeding density of 0.5 × 10^6^ cells/mL and cultured for 14 days. On days 3-12, 3-MA was supplemented at 5 mM and performance was compared against control conditions. While process performance remained comparable until day 5, a reduction in viability and IVCD was observed thereafter for 3-MA treated cultures (Figure 3A). On day 14, viability in 3-MA-treated cultures was approximately 70%, compared to 86% in controls. Specific productivity (Qp) increased by 35%, but this was offset by reduced viable cell density (VCD), leading to a marginally lower titer. The LDH activity in the 3-MA-treated culture was 58% lower than in the control suggesting that the decrease in VCD was potentially due to cell growth arrest rather than cell lysis (Figure 3A). We also monitored key apoptotic proteins (CASP3, CASP8, BAX, CYCS or Cytochrome C, PARP1) which are expected to increase in abundance in the cell culture fluid upon cell lysis due to apoptotic cell lysis. We did not observe any significant change in their level in the HCCF. A dose ranging study optimizing the concentration and treatment duration of 3-MA might help mitigate the low viability concern in the future.

**Figure 3:**
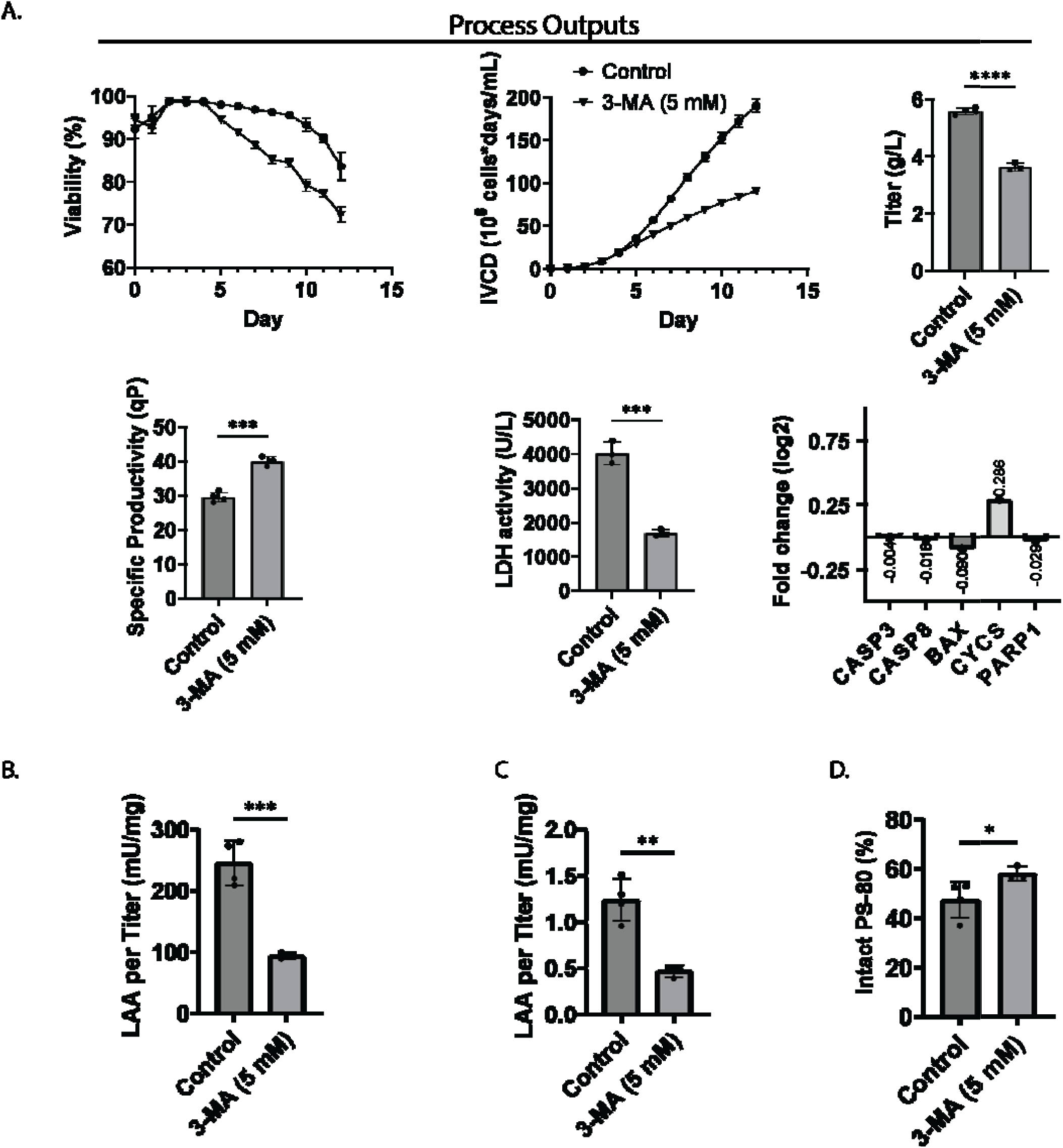
Evaluation of 3-MA effects on cell culture performance and HCP characteristics in fed-batch culture. (A) Fed-batch culture performance, including viability (%), viable cell density (VCD, ×10 cells/mL), titer (g/L), specific productivity (Qp, pg/cell/day), lactate dehydrogenase (LDH) activity (U/L), and log2 fold change (3-MA/Control) of apoptotic protein abundance comparing control and 3-MA (5 mM) conditions. (B) Lipase activity assay (LAA) differences between control and 3-MA conditions, normalized per titer (mU/mg) in HCCF. (C) Lipase activity differences in purified active product (PAP). (D) Intact PS-80 level in PAP after two-week incubation, expressed as a percentage of its level before incubation. Error bars represent the standard error of the mean. Student’s t-test; ns: not significant; *: p < 0.05; **: p < 0.01. *n* _Control_ = 4; *n* _3-MA_ = 3.

Batch culture showed a significant decrease in the abundance of lipases in 3-MA treated cultures. We tested if this difference in abundance translated to reduced lipase activity. In this regard, harvested cell culture supernatant from Day 14 of the fed-batch cultures were analyzed using in-house lipase activity assay. As shown in Figure 3B, cell culture supernatant from 3-MA treated cultures demonstrated a substantial decrease (∼2.6 fold or 62 %) in lipase activity.

During biologics manufacturing processes, once cell culture fluid is harvested, it is processed via a series of chromatography and filtration steps to obtain the final drug substance. At this stage, presence of active lipases or PSDEs can cause the degradation of polysorbate that can impact shelf life of the drug product. We tested if the difference in lipase abundance and activity observed in the cell culture fluid of the 3-MA treated fed batch culture also translate to meaningful differences in PS-80 degradation kinetics after Protein A purification. To assess this, cell culture supernatants from fed batch experiments of 3-MA treated and control cultures was purified using ProA chromatography. Consistent with the significant decrease observed in HCCF, the analyzed ProA pool (PAP) samples of 3-MA treated cultures had a 2.6-fold reduction in lipase activity per mAb (Figure 3C). While lipase activity is often used as a proxy for polysorbate degradation, assay variability and substrate specificity necessitates direct measurement ^31^. In this regard, Protein A pool (PAP) samples were spiked with 0.25 mg/mL PS-80 and staged at 37 °C for 2 weeks. Residual PS-80 concentration was measured to monitor PS-80 degradation kinetics. As shown in Figure 2D, 22 % improvement in PS-80 stability was observed in PAP of 3-MA treated cultures. These findings indicate that 3-MA treatment in fed-batch cultures can selectively reduce PSDE activity and improve PS-80 stability of biologics.

### 3.4 Prolonged autophagy increases specific productivity and afucosylation levels of the mAb

Increased mAb productivity is desirable to improve cost of goods. It has been reported that autophagy influencing compounds such as 3-MA increase specific productivity which is consistent with our data (35 % in our FB harvest) ^14^. Glycosylation levels of biologics can influence protein stability, potency, absorption or clearance ^32^. It has been demonstrated that the absence of fucose in IgG mAbs enhances the antibody-dependent cell mediated cytotoxicity (ADCC) activity ^33^ and antibody-dependent cell-mediated phagocytosis (ADCP) ^34^. Similarly, galactosylation has been shown to enhance CDC activity and degalactosylation reduced CDC activity ^35^. Related studies showed that the difference in the levels of afucosylated glycoforms among REMICADE® (10.0%), INFLECTRA® (6.2%) resulted in a significant difference in the relative FcγR-IIIa receptor binding affinities and subsequent difference in ADCC activity (99.8% vs 50.3%) ^36^. We analyzed the N-glycan profile of the 3-MA treated PAP samples. We observed statistically significant increase of galactosylation as well as afucosylation levels with 3-MA treatment (Figure 4A). Review of the glycosylation data shows that the G0F glycan, which is processed in the Golgi, was significantly reduced in 3-MA-treated samples, while the non-fucosylated G0 species showed an increase (Figure 4B). This suggests a potential reduction in fucosyltransferase activity under 3-MA treatment, contributing to the observed increase in afucosylation. Further experiments are required to confirm this hypothesis. Overall, the reduction in G0F species appears to drive the overall afucosylation increase, highlighting 3-MA’s potential to influence glycosylation in a meaningful way.

**Figure 4:**
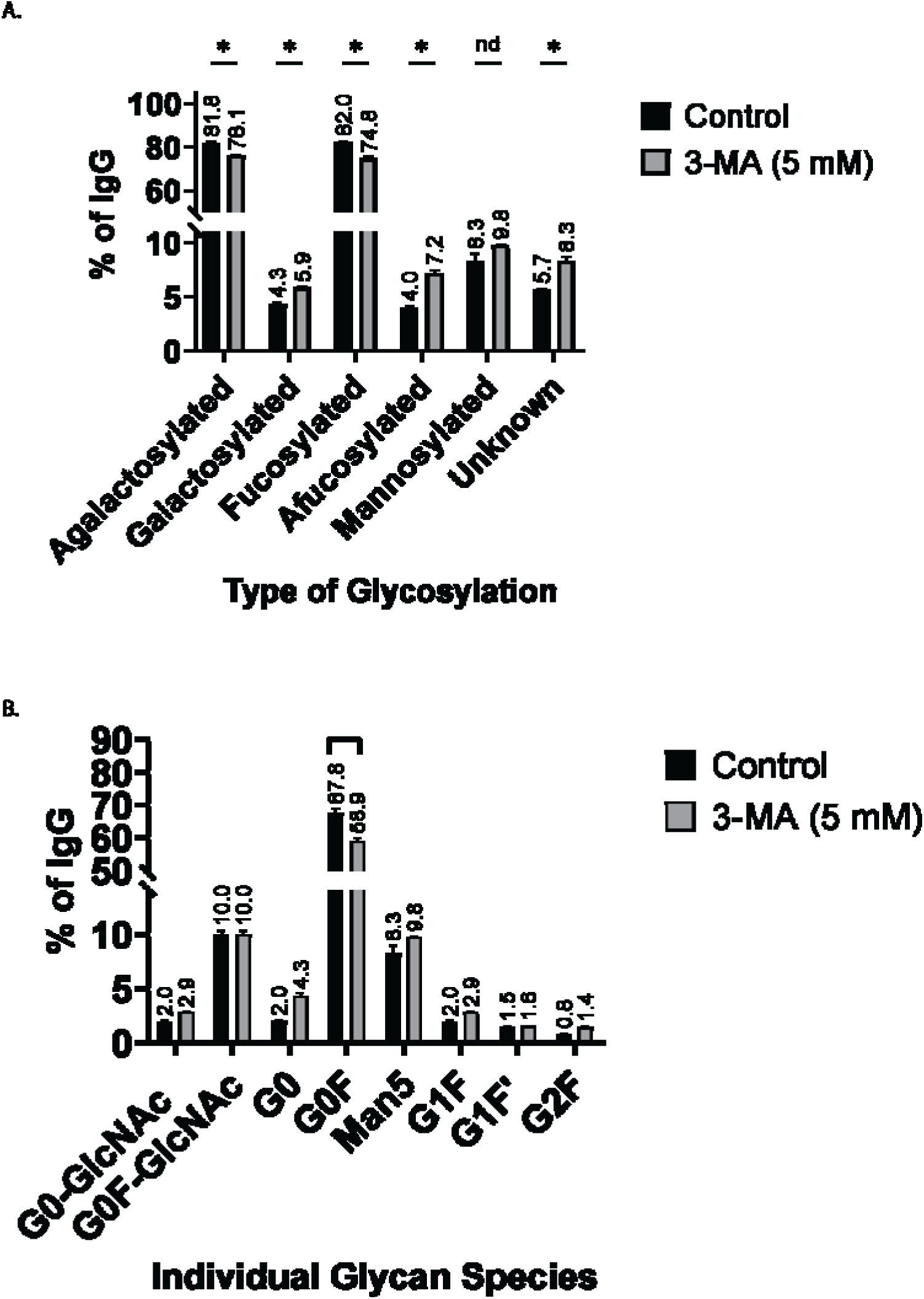
Impact of 3-MA treatment on glycosylation profiles of IgG produced in CHO cells. (A) Distribution of glycosylation types as a percentage of total IgG. Cells were treated with 5 mM 3-MA (gray bars) or left untreated (black bars). (B) Abundance of individual glycan species as a percentage of total IgG for control (black bars) and 3-MA-treated (gray bars) conditions. Asterisks (*) represent statistically significant differences (p < 0.05) and ‘ns’ denotes no statistical difference. Data are presented as mean ± standard deviation of three biological replicates

### 3.5 Autophagy induction measured by LC3B conversion correlates with G1 cell cycle arrest

To gain further insights into the autophagy dynamics induced by 3-MA treatment, we examined its impact on autophagic flux and cell cycle regulation under our experimental conditions. While previous studies suggest that 3-MA promotes autophagic flux in nutrient-rich environments, we sought to confirm this observation in our setup for added reliability. Additionally, we analyzed key cell cycle markers to assess whether 3-MA-mediated autophagy induction is linked to specific cell cycle changes. LC3B-II formation from LC3B-I is widely regarded as a gold-standard marker for autophagy induction. During autophagy, LC3B-I converts to LC3B-II, which integrates into the autophagosomal membrane. The intracellular LC3B-II content reflects autophagosome formation and can be used to monitor autophagic flux.

As shown in Figures 5A (top left) and quantified in 5B (top right), the intracellular LC3B-II (lower western band) to LC3B-I (upper western band) ratio was higher on early culture days (D5 and D7), indicating increased autophagic flux in the presence of 3-MA. To assess autophagy completion, we analyzed p62, a selective autophagy substrate that binds ubiquitinated proteins and is degraded upon autophagosome-lysosome fusion. Reduced p62 levels typically indicate efficient autophagic degradation. As shown in Figures 5A (top left) and quantified in 5B (top right), the intracellular LC3B-II (lower western band) to LC3B-I (upper western band) ratio was higher on early culture days (D5 and D7), indicating increased autophagic flux in the presence of 3-MA. To assess autophagy completion, we analyzed p62, a selective autophagy substrate that binds ubiquitinated proteins and is degraded upon autophagosome-lysosome fusion. Reduced p62 levels typically indicate efficient autophagic degradation ^37, 38^; however, we observed comparable P62 levels in both the conditions, suggesting similar completion of autophagy. autophagy remained active in both conditions, as LAMP1 protein levels were unchanged. LAMP1 plays a crucial role in autophagosome-lysosome fusion, a key step in autophagic flux. In line with the upregulation of autophagy, we observed increased expression of TFEB, the master regulator of autophagy, across culture days (except D10) in 3-MA-treated cultures, consistent with our transcriptomic data. Overall, these findings confirm that 3-MA treatment induces prolonged autophagy, reinforcing its role in modulating autophagic flux in CHO cell cultures.

**Figure 5:**
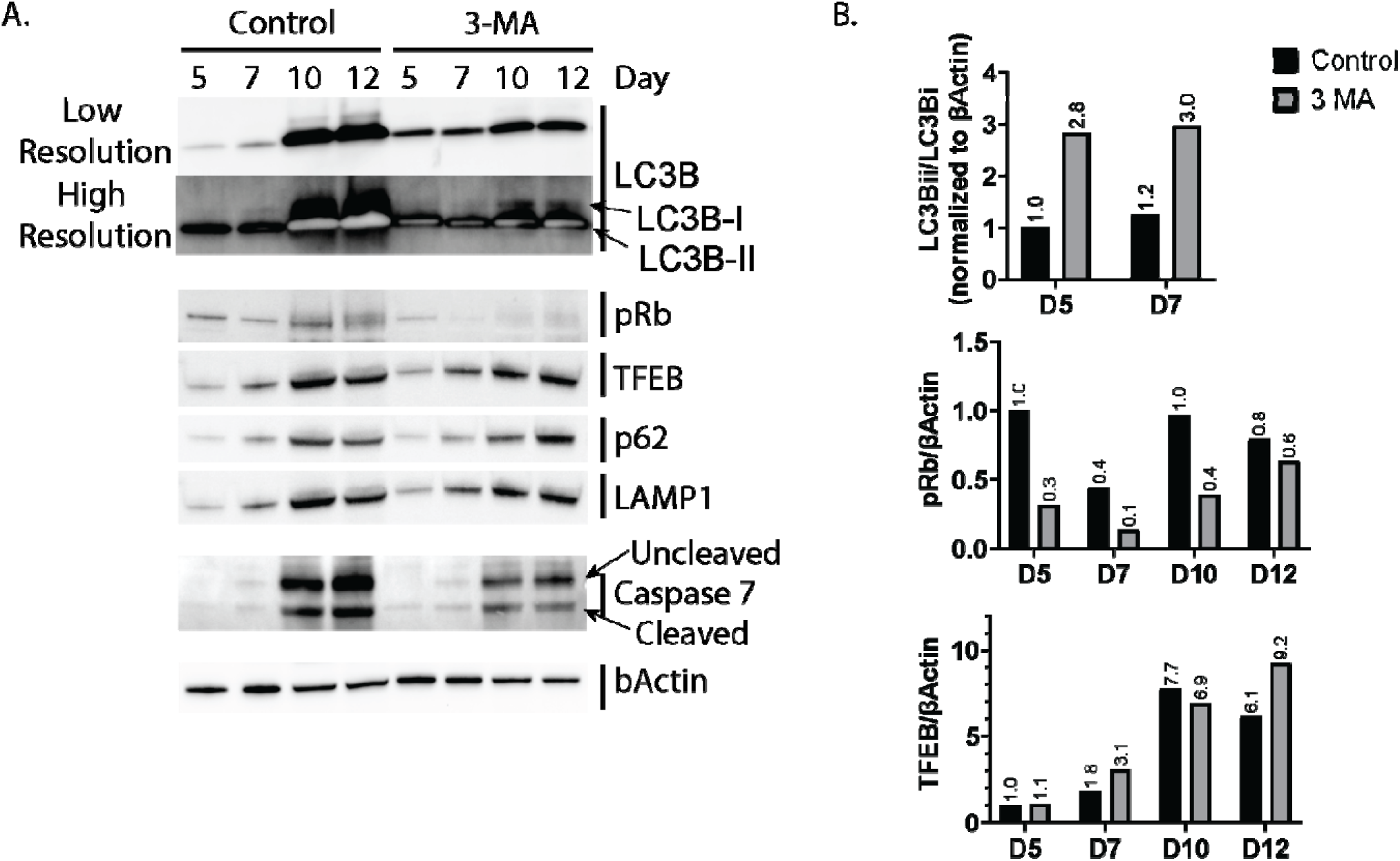
Evaluation of autophagy-related proteins and transcription factors in fed-batch CHO cell culture under 3-MA condition. (A) Representative western blot analysis of LC3B, pRb, TFEB, p62, and LAMP1, Caspase 7 with β-Actin as a loading control, from cell lysates collected on Days 5, 7, 9, and 12. (B) Quantification of selected western blot results normalized to β-Actin. Bars represent mean values, with fold changes labeled.

Our data showed that 3-MA increased specific productivity without much increase in the VCD and cell death (low LDH level). We suspected a cell cycle arrest which could explain this phenomenon. So, we tested to see if there is a cell cycle arrest by monitoring the level of retinoblastoma protein (pRb). This protein controls G1-to-S phase transition with phosphorylation enhancing the cell cycle progression ^39^. We observe that the level of phosphorylated Rb protein is down throughout the culture days in the 3-MA treatment suggesting greater proportion of cells arrested in G0/G1 phase (Figure 5 A, B). These results suggest for the first time that 3-MA initiates autophagy through the engagement of the CLEAR signaling pathway and causes cell cycle arrest which we are reporting for the first time.

## 4. Conclusions

This study highlights the potential of modulating the CLEAR signaling pathway and autophagy to influence lipase activity and polysorbate degrading enzyme (PSDE) levels in CHO cell cultures. We observed upregulation of the CLEAR pathway in Clone B was associated with a reduction in lipase activity and high-risk HCPs, including lipases and PSDEs, as observed in the proteomics analysis. To our surprise, many of the autophagy regulating genes regulated by CLEAR pathway such as BECN1, HIF1, VPS8, etc. were not significantly upregulated in the same clone per our transcriptomics analysis from Day 14. It is possible that on Day 14, the autophagy induction pathway is not active compared to the initial days. In fact, autophagy modulators, such as 3-MA were able to induce autophagy in the earlier days and resulted in reduction of PSDEs, improved product quality, with a significant reduction in lipase activity, increased specific productivity, and enhanced polysorbate stability. Furthermore, the application of 3-MA led to altered glycosylation profiles, including increased afucosylation and galactosylation, which are generally associated with enhanced therapeutic activity. It is possible that these glycosylation changes are a result of G1 phase arrest upon 3-MA treatment. Valeric acid supplementation has been shown to induce G1 arrest along with increased galactosylation and decreased G0F (fucosylated) species ^40^. In addition, specific arrest of CHO cells in G1 phase with CCI (cell cycle inhibitor) compound increased mAb galacosylated species (G1F and G2F) which were attributed to the upregulation of key processing enzymes in glycan trimming and maturation, including GlcNAc beta 1,4 (1,3)galactosyltransferase (B3galt, B4galt), beta-galactosidealpha2,3-sialyltransferase 1 (St3gal), UDP-galactose transporter (Slc35a2) ^41^. Our data indicate that LDH activity, a marker of cell death, is significantly lower in the HCCF of the 3-MA treatment group at the time of harvest, and there is no increased abundance of apoptotic proteins in this group. As a confirmatory test, we measured the caspase-7 cleavage as the marker of apoptosis ^42^. We observed similar levels of caspase-7 cleavage in both control and 3-MA-treated cultures (Figure 5), indicating that 3-MA does not enhance apoptosis. While increased autophagy can sometimes elevate the likelihood of cell death, our data suggest that the reduced viability observed with 3-MA treatment is not driven by apoptosis. Instead, the decrease in viability may result from growth halting due to net ATP depletion arising from accelerated mitophagy exceeding ATP replenishment, increased ATP demand for autophagosome formation and fusion, or the heightened requirement for active proton pumping to maintain lysosomal acidity during degradation ^43^. Additionally, prolonged G1 phase arrest and oxidative stress induced by high cellular productivity could further contribute to reduced viability. Despite this, the upregulation of TFEB and the reduction of lipase activity suggest that 3-MA may provide a viable approach to improving product quality without significant cell toxicity or loss of productivity. These findings offer valuable insights into optimizing biologics production in CHO cells and could guide future efforts in fine-tuning autophagy pathways for better process control.

## Authorship contribution statement

Ansuman Sahoo: Experiment, Data generation, Data Analysis, Visualization, Writing - original draft, Writing - review & editing; Taku Tsukiadate: Experiment, Data generation, Data Analysis, Visualization, Writing - review & editing; Geetanjali Pendyala: Experiment, Data generation; Xuanwen Li: Writing - review & editing, Supervision; Sri R. Madabhushi: Conceptualization, Writing - review & editing, Supervision.

## Acknowledgements

The authors would like to acknowledge the support from colleagues in the MRL Biologics Process Development and Analytical Research & Development group. The authors thank Mike Caruso and Billy Alcaide for media preparation, the in process analytical group for titer and quality measurements, Kaniz Fatema for help with proteomics sample processing, Divya Chandra for help with downstream analysis. The authors would like to thank Richard DJ Chen, and Sanjeev Ahuja for reviewing the manuscript and proving valuable feedback. A.S and T.T. acknowledge support from the MRL Postdoctoral Research Program.

This work was supported by Merck Sharp & Dohme LLC, a subsidiary of Merck & Co., Inc., Rahway, NJ, USA. The authors are employees of Merck Sharp & Dohme LLC, a subsidiary of Merck & Co., Inc., Rahway, NJ, USA, and were responsible for the study design, data collection and analysis, decision to publish, and preparation of the manuscript.

